# Contemporary human H3N2 influenza A viruses require a low threshold of suitable glycan receptors for efficient infection

**DOI:** 10.1101/2022.11.16.516725

**Authors:** Cindy M. Spruit, Igor R. Sweet, Theo Bestebroer, Pascal Lexmond, Boning Qiu, Mirjam J.A. Damen, Ron A. M. Fouchier, Joost Snijder, Sander Herfst, Geert-Jan Boons, Robert P. de Vries

## Abstract

Recent human H3N2 influenza A viruses (IAV) have evolved to employ elongated glycans terminating in α2,6-linked sialic acid as their receptors. These glycans are displayed in low abundancies by cells commonly employed to propagate these viruses (MDCK and hCK), resulting in low or no viral propagation. Here, we examined whether the overexpression of the glycosyltransferases B3GNT2 and B4GALT1, which are responsible for the elongation of poly-N-acetyllactosamines (LacNAc), would result in improved A/H3N2 propagation. Stable overexpression of B3GNT2 and B4GALT1 in MDCK and hCK cells was achieved by lentiviral integration and subsequent antibiotic selection and confirmed by qPCR and protein mass spectrometry experiments. Flow cytometry and glycan mass spectrometry experiments using the B3GNT2 and/or B4GALT1 knock-in cells demonstrated increased binding of viral hemagglutinins and the presence of a larger number of LacNAc repeating units, especially on hCK-B3GNT2 cells. An increase in the number of glycan receptors did, however, not result in a greater infection efficiency of recent human H3N2 viruses. Based on these results, we propose that H3N2 IAVs require a low number of suitable glycan receptors to infect cells and that an increase in the glycan receptor display above this threshold does not result in improved infection efficiency.

## Introduction

Influenza A viruses (IAV) of the H3N2 subtype cause seasonal epidemics, leading to illness, hospitalizations, and deaths in humans [1]. The crucial first step of infection is the binding of the viral hemagglutinin (HA) to a receptor on a cell, which are glycans terminating in α2,6-linked sialic acids (SIA) for human IAVs [2, 3]. H3N2 viruses have been circulating in the human population since 1968 and due to continuous immune evasion, antigenic drift of the surface proteins of IAVs takes place. This antigenic drift of H3N2 viruses has changed receptor specificities [4] and recent H3N2 viruses bind to longer glycans having multiple consecutive oligo-N-acetyllactosamine (LacNAc) moieties terminating in an α2,6-linked SIA [2, 4-11]. This specificity is most pronounced for H3N2 viruses of subclade 3C.2a, which require at least three subsequent LacNAc repeating units for binding [12].

These altered receptor specificities make it difficult to isolate and propagate H3N2 viruses, greatly hampering the further study of these viruses [8, 13-16]. Even when virus isolation is successful, viruses may have acquired adaptive mutations in the receptor binding site of HA, especially when isolated in eggs instead of MDCK (Madin-Darby Canine Kidney) cells [13, 15, 17-20]. MDCK cells have previously been modified to produce more α2,6-linked SIAs by the overexpression of the enzyme ST6GAL1, resulting in MDCK-SIAT1 [21] and MDCK-AX4 [22] cells. These cells enabled the isolation of H3N2 viruses, especially of the 3C.2a and 3C.3a subclades, and resulted in higher titers of viral stocks [16, 23]. To allow the isolation of further evolved contemporary H3N2 viruses, with higher titers and fewer mutations, MDCK cells were further modified to eliminate α2,3-linked SIAs while also overexpressing α2,6-linked SIAs, resulting in “humanized” MDCK (hCK) cells [15].

Analysis of the *N*-glycans of MDCK, MDCK-SIAT1, and hCK cells indicated a low abundance of glycans with at least three successive LacNAc repeating units terminating in an α2,6-linked SIA [24]. The enzyme beta-1,3-N-acetylglucosaminyltransferase (B3GNT2) is responsible for the addition of N-acetylglucosamine to glycans, while the galactose is transferred to the glycan by the enzyme beta-1,4-galactosyltransferase 1 (B4GALT1). Previously, we successfully used these two enzymes to elongate LacNAc repeating units both in chemoenzymatic synthesis [25, 26] and on biological membrane surfaces of erythrocytes [12].

Here, we genetically engineered MDCK and hCK cells to overexpress B3GNT2 and B4GALT1 and demonstrated that this resulted in a higher relative abundance of *N*-glycans having elongated LacNAc moieties. Surprisingly, although the B3GNT2/B4GALT1 knock-in cells exhibited elevated binding of recent H3 HAs, the overexpression did not lead to improved virus isolation and infection efficiency. Several studies have indicated that a higher display of appropriate receptors leads to increased infectivity [15, 21, 22, 27], while others indicated that only low amounts of receptors are required for infection [28-30]. Based on our studies, we concluded that above a required threshold, a greater number of suitable glycan receptors for H3N2 IAVs does not result in increased infection efficiency.

## Results

### Generation of stable B3GNT2 and B4GALT1 knock-in MDCK and hCK cell lines

Rearrangement of the sialyltransferase expression in hCK cells supported increased replication of many human H3N2 viruses [15]. However, only small quantities of glycans with multiple LacNAc repeating units appeared to be present on both MDCK and hCK cells [24]. Recently, we and others have shown that poly-LacNAc containing *N*-glycans are critical for the binding of contemporary H3N2 viruses [8, 12]. Therefore, we used the glycosyltransferases B3GNT2 and B4GALT1 to increase the biosynthesis of LacNAc repeating units to produce extended *N*-glycans [12, 25, 26, 31]. We hypothesized that the overexpression of B3GNT2 and/or B4GALT1 in MDCK and hCK cells would produce appropriate glycan receptors for recent H3N2 (subclade 3C.2a) IAVs.

To accomplish the overexpression of these genes in MDCK and hCK cells, lentiviral transfer plasmids encoding the human *B3GNT2* and/or *B4GALT1* genes, together with the *Hygromycin B resistance* gene, were constructed. The genes were expressed from one human EF-1α promoter [32] and separated by P2A (and for double glycosyltransferase knock-ins also T2A) self-cleaving peptides. Lentiviruses were produced with a transfer plasmid and packaging plasmids, after which the viruses were used to transduce MDCK and hCK cells (Fig. 1A). Cells in which the genes were inserted in the genome were selected with Hygromycin B.

**Fig 1.**
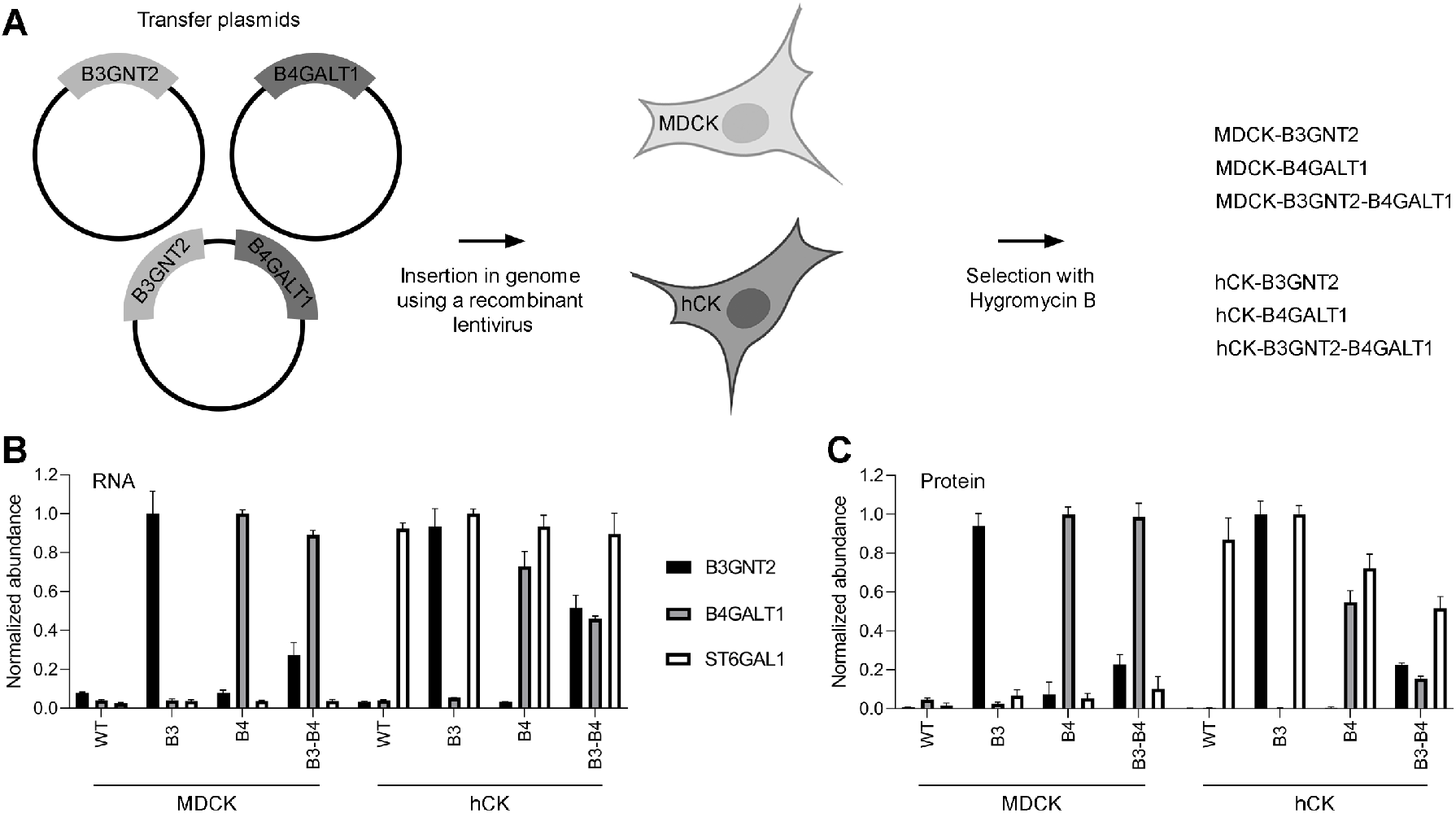
Construction of MDCK and hCK cells that overexpress B3GNT2 and/or B4GALT1. (**A**) MDCK and hCK cells were modified with recombinant lentiviruses containing transfer plasmids for the insertion of the *B3GNT2* and/or *B4GALT1* gene and the Hygromycin B resistance gene. The knock-in cells were selected with 300 µg/ml Hygromycin B. (**B**) RT-qPCR was performed with primers that anneal to both the human and dog *B3GNT2, B4GALT1*, or *ST6GAL1* genes. The values relative to the dog *GAPDH* gene were used, which were then normalized to the highest value of each gene. Mean and SD (n=3) are shown. (**C**) Mass spectrometry of the B3GNT2, B4GALT1, and ST6GAL1 proteins. Only peptides unique to human proteins were selected. All samples were normalized against tubulin beta and then normalized to the highest value of each protein. Mean and SD (n=3) are shown.

Stable overexpression of *B3GNT2* and *B4GALT1* was confirmed by RT-qPCR analysis on isolated cellular RNA. Primers for the *B3GNT2, B4GALT1*, and *ST6GAL1* genes were used, and the obtained values were normalized to the reference gene *GAPDH* (Fig. 1B). Overexpression of the control gene *ST6GAL1* was clearly shown in hCK but not MDCK cells. Furthermore, the overexpression of *B3GNT2* and *B4GALT1* was present in all cell lines in which these knock-ins were made. It should be noted that expression levels in the double knock-in cell lines showed lower expression of the glycosyltransferases, especially for *B3GNT2* in MDCK-B3GNT2-B4GALT1 cells.

Thereafter, the protein levels of B3GNT2, B4GALT1, and ST6GAL1 in cell lysates were measured using proteomic experiments based on liquid chromatography coupled to tandem mass spectrometry, using label-free quantitation relative to tubulin beta expression (Fig. 1C). Only peptides unique for the human B3GNT2, B4GALT1, and ST6GAL1 were selected. The proteomic data is comparable to the RT-qPCR data since elevated protein levels in the cell lines with knock-ins were observed. Collectively, the data showed that the stable overexpression of B3GNT2 and B4GALT1 in MDCK and hCK was successful.

### Flow cytometric characterization of B3GNT2 and B4GALT1 knock-in cells with plant and viral lectins

Next attention was focused on whether the overexpression of B3GNT2 and/or B4GALT1 led to a display of a higher number of LacNAc repeating units on *N*-glycans. The glycans on the cell surface were first characterized using flow cytometry with standard lectins. An alive, single-cell population was selected using a standard gating strategy, and mean fluorescence intensities were calculated over the cell population (Fig. 2).

**Fig 2.**
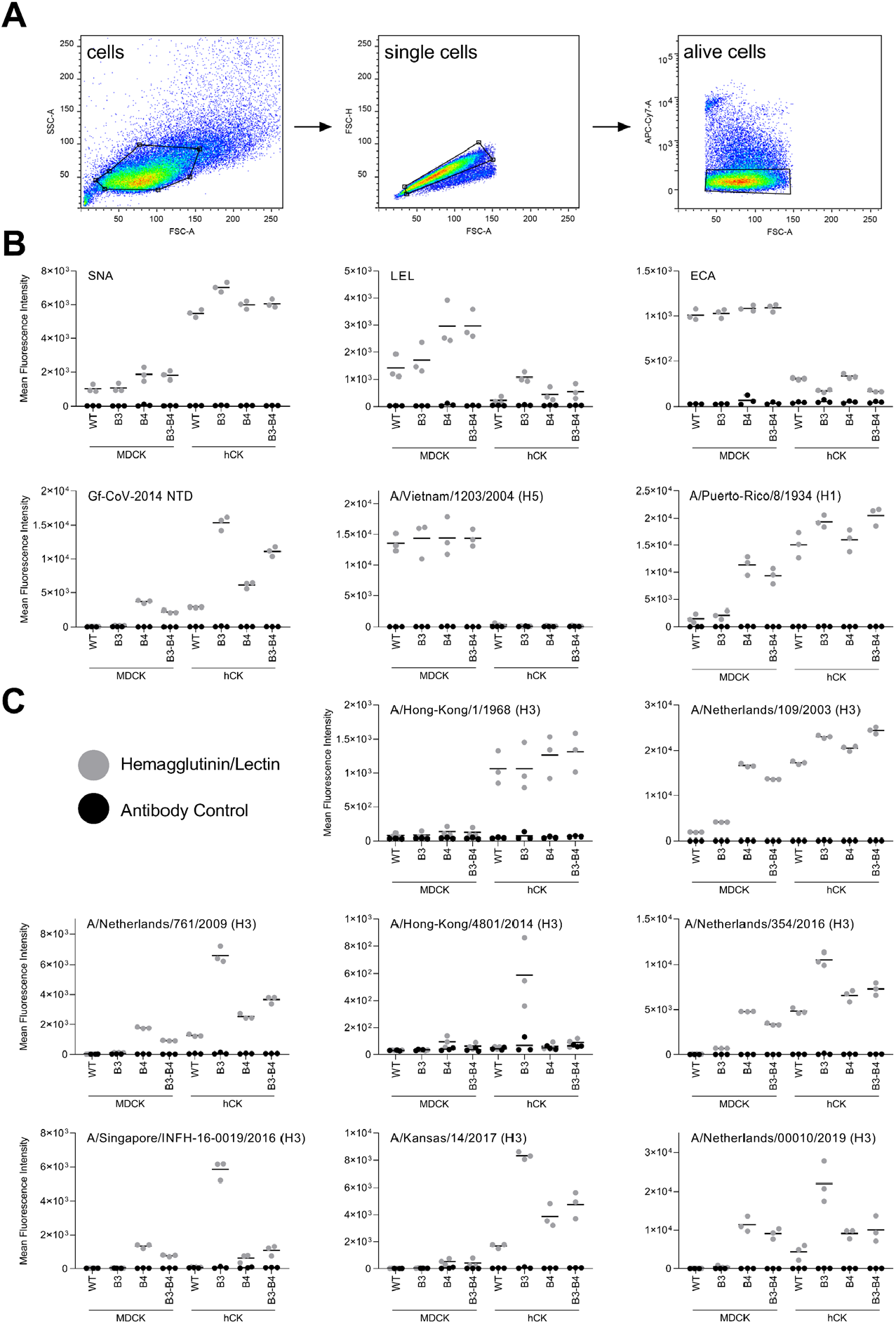
Flow cytometric characterization of B3GNT2/B4GALT1 knock-in MDCK and hCK cells. (**A**) The gating strategy that was used to select single, alive cells. (**B**) Flow cytometry measurements with lectins SNA (*Sambucus nigra* agglutinin, recognizes α2,6-SIA), LEL (*Lycopersicon esculentum* lectin, recognizes elongated glycans), and ECA (*Erythrina cristagalli* lectin, recognizes glycans without SIA) were performed. Furthermore, Gf-CoV-2014 NTD was used to detect elongated glycans, H5 HA of A/Vietnam/1203/2004 was used to detect α2,3-SIA, and H1 HA from A/Puerto-Rico/8/1934 was used as a standard influenza virus. Triplicate measurements were performed, of which the mean and all individual measurements are displayed. (**C**) A diverse set of H3 HAs was used to characterize the cells. Triplicate measurements were performed, of which the mean and all individual measurements are displayed. Titration curves of A/Hong-Kong/1/1968, A/Netherlands/109/2003, and A/Singapore/INFH-16-0019/2016 are shown in Fig. S2. Flow cytometric experiments with neuraminidase treatment of the cells are shown in Fig. S3

*Sambucus nigra* agglutinin (SNA) [33] was used to detect α2,6-linked SIAs, which are present in higher quantities on hCK cells than MDCK cells due to the overexpression of ST6GAL1. The B3GNT2 and/or B4GALT1 knock-ins did not cause substantial differences in α2,6-linked SIA display, which is understandable since we did not interfere with the sialyltransferases. *Lycopersicon esculentum* lectin (LEL) recognizes elongated glycans [34] and we observed that glycans capped with α2,6-linked SIAs need at least four consecutive LacNAc repeating units to be recognized, while glycans capped with α2,3-linked SIAs are recognized when presented on two successive LacNAc repeating units (Fig. S1), which explains the lower signal for all hCK cells in general. A substantial increase in the binding of LEL to MDCK-B4GALT1 and MDCK-B3GNT2-B4GALT1 compared to WT MDCK cells was observed, indicating that the LacNAc repeating units on MDCK cells are indeed elongated by the overexpression of mainly B4GALT1. Moreover, we observed an increase in LEL signal in the hCK-B3GNT2 cells compared to the hCK WT cells, indicating that the overexpression of B3GNT2 resulted in the formation of longer glycans on hCK cells. *Erythrina cristagalli* lectin (ECA) recognizes terminal galactose, and thus glycans lacking SIA capping [35]. The results using ECA indicated that all MDCK cells have a larger proportion of non-sialylated glycans compared to all hCK cells, which agrees with the overexpression of ST6GAL1 in all hCK cells. No major differences in the amount of non-sialylated glycans between WT and B3GNT2 and/or B4GALT1 knock-in cells were observed.

In addition to commonly employed plant lectins, viral proteins were used to examine the glycans displayed on the cells. The N-terminal domain of γCoV/AvCoV/guinea fowl/France/14032/2014 (Gf-CoV-2014 NTD) is known to bind elongated glycans [36]. MDCK-B4GALT1 and MDCK-B3GNT2-B4GALT1 cells showed an increased Gf-CoV-2014 NTD signal compared to MDCK WT cells. Furthermore, hCK WT cells appeared to have a higher number of LacNAc repeating units on glycans than MDCK WT cells. The hCK-B3GNT2 cells, and the hCK-B4GALT1 and hCK-B3GNT2-B4GALT1 cells to a lesser extent, showed a substantial increase in Gf-CoV-2014 NTD binding compared to the hCK WT cells, indicating the presence of additional LacNAc repeating units on glycans. The HA of A/Vietnam/1203/2004 H5 (H5VN) is commonly used to probe the presence of α2,3-linked SIAs [35, 37, 38]. We observed a much lower amount of α2,3-linked SIAs in all hCK cells compared to all MDCK cells, which is in agreement with the knock-outs of all β-galactoside α-2,3 sialyltransferases that were made in the hCK cells previously [15]. The B3GNT2/B4GALT1 knock-ins did not alter the α2,3-linked SIA content. The HA of the human IAV A/Puerto-Rico/8/1934 (PR8) H1 binds α2,6-linked SIAs [39] and showed increased binding to all hCK cells compared to all MDCK cells, which is related to the overexpression of ST6GAL1 in hCK cells [15]. Increased binding of PR8 to MDCK-B4GALT1 and MDCK-B3GNT2-B4GALT1 compared to MDCK WT cells was also observed, which deviates from the results obtained with SNA, with which no increase in α2,6-linked SIAs was shown.

### hCK-B3GNT2 cells are preferentially bound by contemporary H3 HAs

After initial characterization, an array of human H3 HAs was used for flow cytometric binding studies with B3GNT2 and B4GALT1 knock-in cells (Fig. 2C). To cover a broad scope of receptor binding specificities, HAs from viruses from different years (1968-2019) and (sub)clades were chosen. Three HAs from the 3C.2a subclade (A/Singapore/INFH-16-0019/2016, A/Netherlands/00010/2019, and A/Hong-Kong/4801/2014) were chosen to assess the presence of glycans with elongated LacNAc structures [12] on the MDCK and hCK WT and B3GNT2/B4GALT1 knock-in cells.

These human H3 HAs prefer binding to α2,6-linked SIAs over α2,3-linked SIAs and therefore, in general, more binding is observed to hCK WT cells than to MDCK WT cells. The HA of A/Hong-Kong/1/1968 does not require multiple consecutive LacNAc repeating units for binding, but it does show a strong preference for glycans with three or four consecutive LacNAc repeating units compared to glycans with only one or two repeating units [8]. The preference of A/Hong-Kong/1/1968 for longer glycans is however not observed in our flow cytometry experiments, since the B3GNT2 and B4GALT1 knock-in cells did not show increased binding.

The human H3N2 IAVs A/Netherlands/109/2003 and A/Netherlands/761/2009 were previously shown to bind glycans with both two and three, but not one, consecutive LacNAc repeating units [12]. Increased binding of A/Netherlands/109/2003 to MDCK-B4GALT1, MDCK-B3GNT2-B4GALT1, and all hCK cells compared to MDCK WT cells was observed. Interestingly, the MDCK-B4GALT1 and MDCK-B3GNT2-B4GALT1 cells often showed comparable or higher binding to the recent H3 HAs than the hCK WT cells, while low levels of α2,6-linked SIAs are present on all MDCK cells. Similar binding patterns were observed for A/Netherlands/761/2009, although the binding to hCK-B3GNT2 cells was much more pronounced.

Contemporary H3 IAVs are known to bind to glycans with multiple consecutive LacNAc repeating units [2, 4-12]. This binding specificity is most pronounced for H3N2 viruses of subclade 3C.2a, which require at least three subsequent LacNAc repeating units for binding [12]. Increased binding to MDCK-B4GALT1, MDCK-B3GNT2-B4GALT1, and all hCK cell lines compared to MDCK WT cells was observed for the recent H3 HAs (2014-2019) A/Netherlands/354/2016, A/Singapore/INFH-16-0019/2016 (subclade 3C.2a), A/Kansas/14/2017 (3C.3a), and A/Netherlands/00010/2019 (subclade 3C.2a). The hCK-B3GNT2 cells that were already indicated to have the longest glycans by LEL and Gf-CoV-2014 NTD, also showed a substantial increase in binding of these recent H3 HAs compared to all other cell lines investigated. The strong binding to hCK-B3GNT2 cells of A/Hong-Kong/4801/2014 (subclade 3C.2a) was even more obvious, as other cell lines appear to be barely bound to this HA.

### Lectin binding to cells is concentration and sialic acid-dependent

To investigate whether the binding of the H3 HAs in the flow cytometry experiments was indeed specific for SIAs, we performed experiments with neuraminidase-treated cells (Fig. S2). As controls for the removal of α2,3-linked or α2,6-linked SIAs, A/Vietnam/1203/2004 [35, 37, 38] and SNA [33] were used. These lectins showed a substantial decrease in the binding signal after neuraminidase treatment. When testing two H3 HAs with well-defined binding specificities, A/Netherlands/109/2003 and A/Netherlands/761/2009 [12], similar results were obtained, indicating that the binding of H3 HAs was indeed SIA-dependent

When examining the binding of the HAs of human H3N2 viruses to cells in Fig. 2C, there appeared to be no binding to the MDCK WT cells at all for A/Hong-Kong/1/1968, A/Netherlands/761/2009, A/Hong-Kong/4801/2014, A/Netherlands/354/2016, A/Singapore/INFH-16-0019/2016, A/Kansas/14/2017, and A/Netherlands/00010/2019, even though it is possible to propagate these viruses in MDCK WT cells. To investigate whether binding to MDCK cells occurred at all, a titration with H3 HAs was performed. As a positive control, A/Netherlands/109/2003 was used since binding was observed in Fig. 2C. Furthermore, the HAs of the well-defined A/Hong-Kong/1/1968 [8] and the subclade 3C.2a virus A/Singapore/INFH-16-0019/2016 were used. The titration indicated that there is indeed binding to MDCK WT cells but to a much lesser extent than to hCK WT (or hCK-B3GNT2 cells) (Fig. S3). From the flow cytometric experiments, it appeared that the hCK-B3GNT2 cells present the highest number of LacNAc repeating units compared to the other cell lines investigated.

### Elongated glycans are detected on hCK-B3GNT2 and hCK-B4GALT1 cells

Since *N*-glycans are the most relevant receptors on cells for IAV [40], we further investigated the *N*-glycans of WT and B3GNT2/B4GALT1 knock-in MDCK and hCK by mass spectrometry of released *N*-glycans. Compared to MDCK WT cells, all seven other cell lines showed a large reduction in the relative abundance of high-mannose glycans (Fig. S4A), to which IAV does not bind. This increase in the relative abundance of complex and hybrid *N*-glycans may partially explain the improved binding phenotype of the H3 HAs to cell lines different than MDCK WT as observed in the flow cytometry experiments (Fig. 2C).

From the flow cytometry experiments, the hCK-B3GNT2 cells were expected to have the highest number of LacNAc repeating units compared to the other seven cell lines (Fig. 2C). The data obtained from the glycan mass spectrometry experiments indeed showed a higher relative abundance of elongated glycans with more than four LacNAc repeating units in hCK-B3GNT2 compared to hCK WT cells (Fig. 3A). Since we were unable to determine the exact structure of the glycans, glycans with at least four LacNAc repeating units were considered as potential receptors for contemporary H3N2 IAVs, since one of the LacNAc repeating units is often present on the other arm [12] and three consecutive LacNAc repeating units are required.

**Fig 3.**
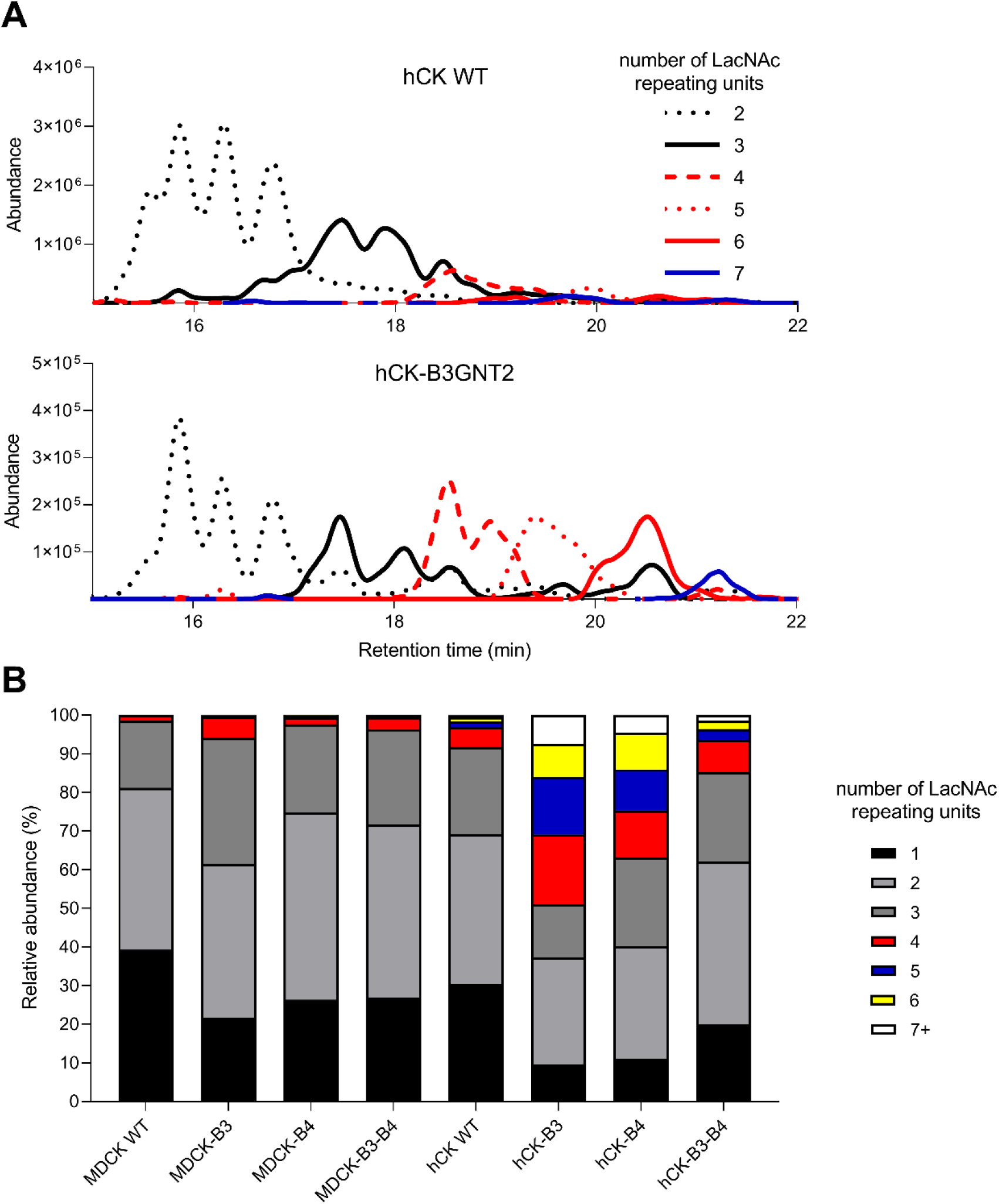
*N*-glycan analysis of WT and B3GNT2/B4GALT1 knock-in MDCK and hCK cells using mass spectrometry. The *N*-glycans from WT and B3GNT2/B4GALT1 knock-in MDCK and hCK cells were measured using mass spectrometry. (**A**) Chromatograms of hCK WT and hCK-B3GNT2 cells were constructed for the glycans with at least two and at most seven LacNAc repeating units. The extracted-ion-counts for the ten most abundant glycan features per LacNAc repeating unit group were summed to yield a chromatogram. (**B**) The *N*-glycans with at least one LacNAc repeating unit were analyzed for the number of LacNAc repeating units present and the relative abundance was calculated. Further analysis is presented in Figure S4. Full glycan feature lists for each cell line are presented in Table S1-8.

The *N*-glycans with at least one LacNAc repeating unit were further analyzed to determine the relative abundance of glycans with a different number of LacNAc repeating units in the eight different cell lines (Fig. 3B). Knock-in cell lines MDCK-B4GALT1 and MDCK-B3GNT2-B4GALT1 did not show an increase in the number of LacNAc repeating units compared to MDCK WT cells. In MDCK-B3GNT2 and hCK WT cells, the relative increase of glycans with four LacNAc repeating units was a few percent compared to MDCK WT cells. The relative abundance of elongated glycans was even higher in hCK-B3GNT2-B4GALT1 cells. Surprisingly, hCK-B4GALT1 showed a substantial increase in the relative abundance of glycans with a higher number of LacNAc repeating units, up to even nine LacNAcs (Table S7), which was not expected from the results of the flow cytometry experiments. The highest increase in the relative abundance of elongated glycans was observed in hCK-B3GNT2 cells, which agrees with the flow cytometric results.

For binding of contemporary H3N2 IAVs, *N*-glycans with at least three consecutive LacNAc repeating units should also be capped with α2,6-linked SIAs. While we were unable to determine the SIA linkage, we analyzed the percentage of sialylation of the glycans with at least one LacNAc repeating unit (Fig. S4B). In general, 75-96% of these *N*-glycans were sialylated, except for the glycans of MDCK-B3GNT2 cells (38% sialylated), which correlated with the low binding of H3 HAs as observed using flow cytometry (Fig. 2C). Whereas non-sialylated glycans occurred in all groups of LacNAc lengths in MDCK WT and B3GNT2/B4GALT1 knock-in cells, all glycans with at least four LacNAc repeating units (except for 4 glycans in total) on all hCK cell lines were sialylated (Table S1-8).

### Sugar nucleotides are not a limiting factor in the biosynthesis of poly-LacNAc structures

Changes in sugar nucleotide levels have been observed in cells after overexpression of B3GNT2 and B4GALT1 [31], which may be a limiting factor in the elongation of glycans. Therefore, the concentrations of sugar nucleotides in the cell lysates of MDCK WT, hCK WT, and hCK-B3GNT2 cells were measured by mass spectrometry (Fig. 4, with details in Fig. S5) [41]. The overexpression of ST6GAL1 in hCK WT and hCK-B3GNT2 cells compared to MDCK WT cells resulted in elevated levels of sialylation and thereby lower levels of available CMP-Neu5Ac, which was also demonstrated in the sugar nucleotide analysis. No other major differences or depletions of sugar nucleotides were observed in any of the cell lines. Most importantly, the sugar nucleotides that are required for the biosynthesis of LacNAc repeating units (UDP-galactose and UDP-HexNAc) were not depleted in any of the cell lines, thus sugar nucleotide availability is likely not a limiting factor for the elongation of glycans.

**Fig 4.**
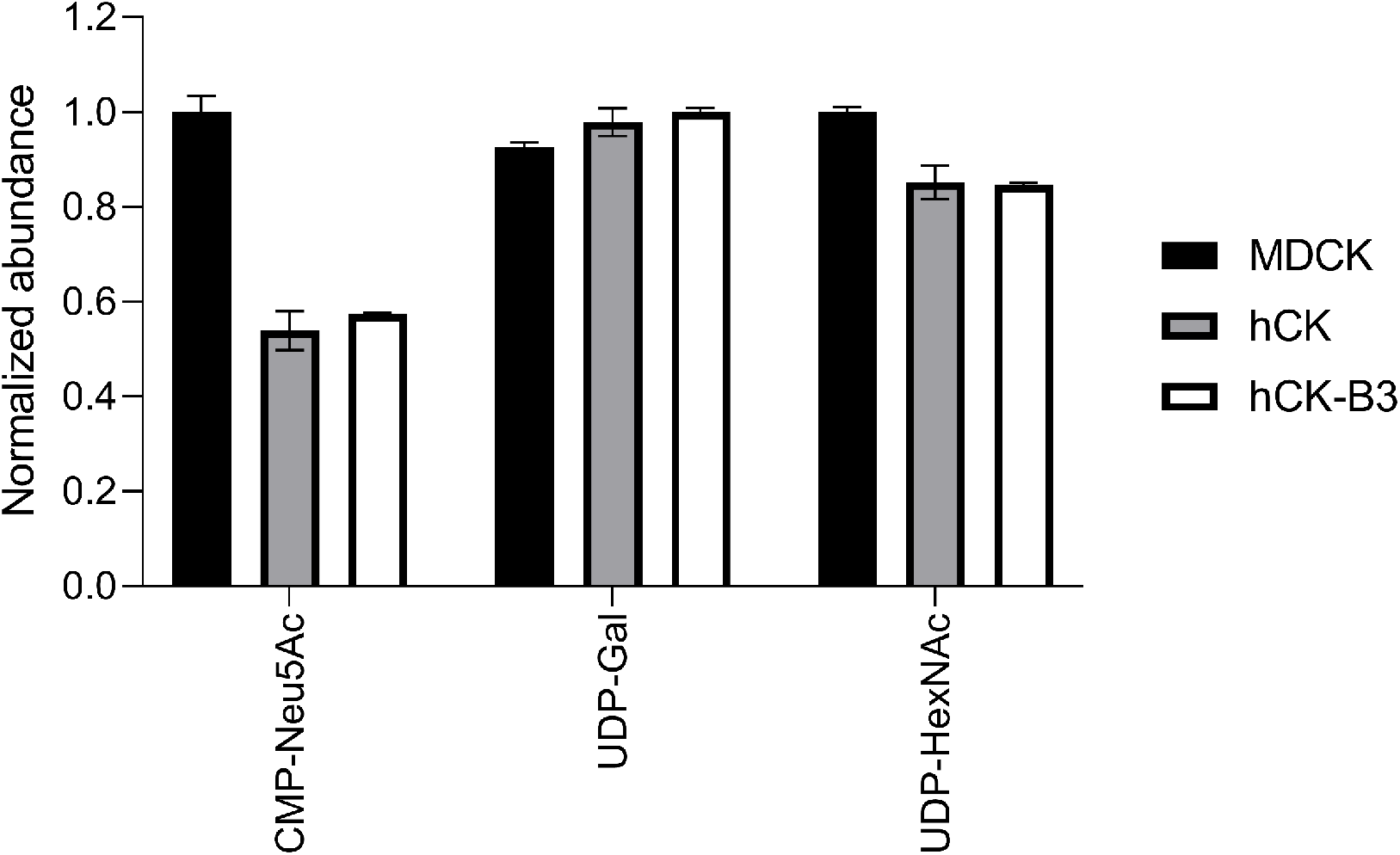
Sugar nucleotide analysis of MDCK, hCK, and hCK-B3GNT2 cells. The sugar nucleotides in the lysate of MDCK, hCK, and hCK-B3GNT2 cells were analyzed by mass spectrometry (n=2). The normalized abundance of CMP-Neu5Ac, UDP-Gal, and UDP-HexNAc are shown. Normalization was performed on the cell line with the highest amount of each sugar nucleotide. Detailed information about all measured sugar nucleotides is presented in Fig. S4.

### Improved binding of hemagglutinins to cells does ensure higher virus titers

MDCK WT, hCK WT, and B3GNT2 and/or B4GALT1 knock-in cells were inoculated with H3N2 viruses to investigate whether higher titers could be obtained in the knock-in cells. Four control viruses (H3N2 from 2003, H1N1, and influenza B, Fig. 5A) and eight recent (2017-2019) H3N2 viruses from the 3C.2a (Fig. 5B) and 3C.3a (Fig. 5C) subclades were used for inoculation. For the H3N2 virus from 2003, the H1N1 virus, and the influenza B viruses, no substantial difference was observed between the virus titers obtained in MDCK, hCK, or B3GNT2/B4GALT1 knock-in cells (Fig. 5A).

**Fig 5.**
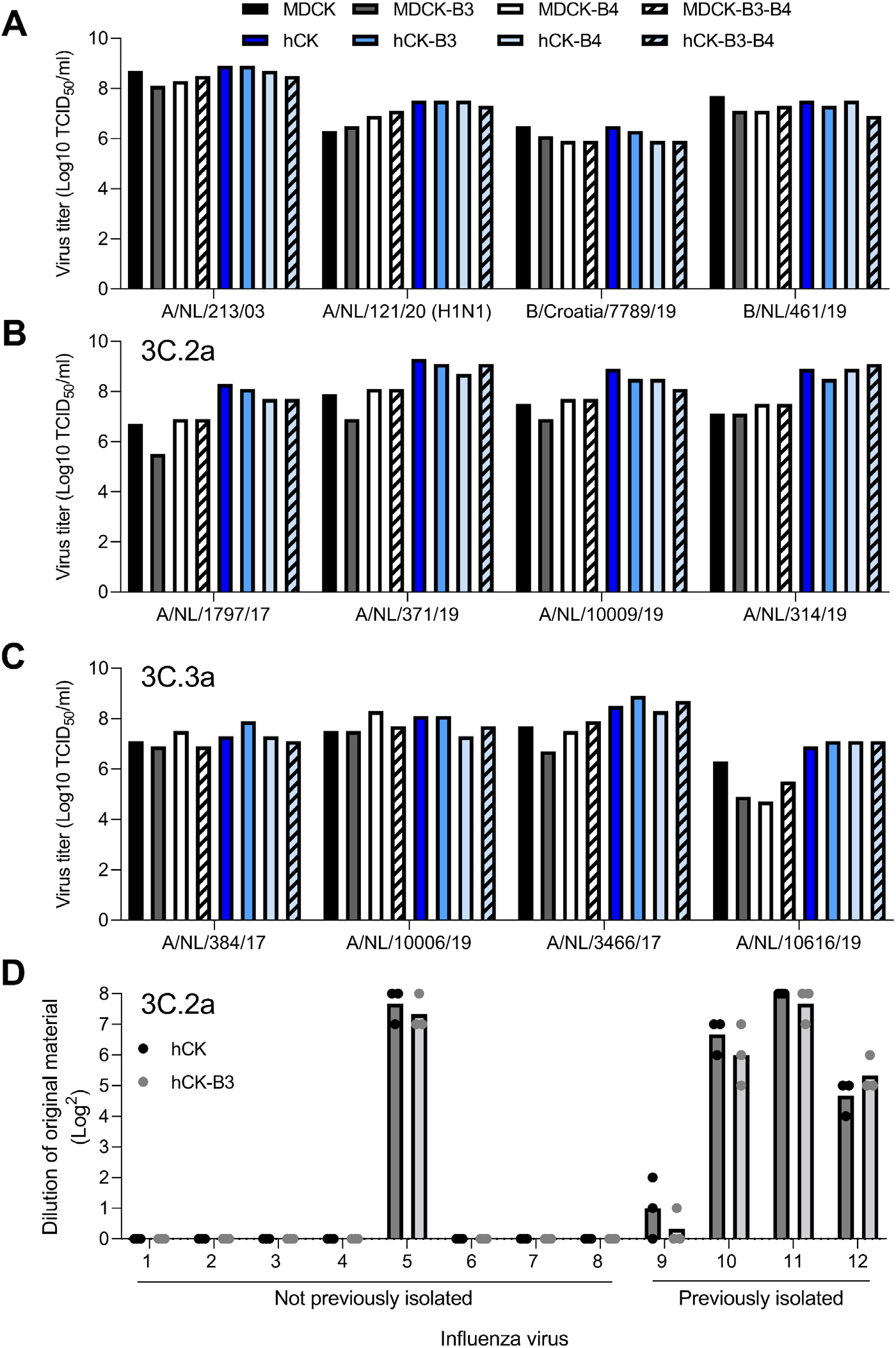
Influenza virus inoculation of B3GNT2 and B4GALT1 knock-in MDCK and hCK cells. End-point titrations with four control viruses and eight recent H3N2 IAVs (details in Table 3) were performed, of which (**A**) four control viruses, (**B**) four 3C.2a viruses, and (**C**) four 3C.3a viruses. Infectious titers were determined either using a hemagglutination assay (**A**) or a nucleoprotein staining (**B, C**) when a hemagglutination assay was not possible. (**D**) An infection study using hCK and hCK-B3GNT2 cells with a twofold dilution of twelve H3N2 IAVs from the 3C.2a clade (details in Table 4) which could previously either not be isolated in hCK cells (#1-8) or could be isolated in hCK cells (#9-12) was performed. Infection was assessed by the presence of cytopathic effects. Individual and mean values are shown.

Recent 3C.2a viruses are known to only bind glycans having at least three consecutive LacNAc repeating units, while recent 3C.3a viruses also bind glycans with two consecutive LacNAc repeating units [12]. For the 3C.2a viruses, a considerable difference was visible between the titers in MDCK WT and hCK WT cells (Fig. 5B), which correlates with the increased binding as observed in the flow cytometric experiments (Fig. 2). Surprisingly, no substantial difference between the titers in hCK WT and hCK-B3GNT2 cells was observed, while the glycans on the latter cell line were extended as observed in the flow cytometry (Fig. 2) and glycan mass spectrometry experiments (Fig. 3). Furthermore, no difference was observed in the titers for the 3C.3a viruses (Fig. 5C), not even between MDCK and hCK cells. Therefore, we concluded that additional binding does not necessarily lead to a higher infection efficiency.

### Isolation of influenza viruses in hCK-B3GNT2 cells is not improved as compared to hCK WT cells

While additional H3N2 viruses can be isolated in hCK cells as compared to MDCK-SIAT1 and MDCK cells [15], we noticed that some H3N2 viruses could still not be isolated in hCK cells. To investigate whether cell lines with longer glycans (hCK-B3GNT2 cells) would facilitate the isolation of additional viruses, we attempted to isolate twelve H3N2 viruses (clade 3C.2a) from original patient material in hCK and hCK-B3GNT2 cells (Fig. 5D). Eight of those viruses could not be isolated previously since they did not replicate in hCK cells. All viruses that were previously isolated in hCK cells were again successfully isolated. However, the use of hCK-B3GNT2 cells did not result in more efficient isolation of those viruses. One of the viruses that was not isolated previously (A/Netherlands/173/2019) could now be isolated in both hCK and hCK-B3GNT2 cells. None of the other viruses that could not be isolated previously could now be isolated in either hCK or hCK-B3GNT2. Therefore, a higher number of extended glycans did not improve the isolation of H3N2 IAVs from the 3C.2a clade.

## Discussion

Although we elongated the LacNAc repeating units on the glycans of MDCK and hCK cells, it did not result in a higher infection efficiency of recent H3N2 IAVs. It has been reported that a low but critical threshold of high-affinity receptors is required for infection [29, 30], though binding and infection are further assisted by the presence of high-abundance low-affinity receptors [28]. This implies that increased HA binding will lead to enhanced entry efficiency. It is therefore counterintuitive that presenting preferred ligands in copious amounts does not lead to increased infection. Strikingly, here we demonstrated that increased HA binding to cells does not necessarily result in more efficient infection.

On the other hand, several studies have indicated that increasing the number of preferred receptors will increase the infection efficiency of IAVs [15, 21, 22, 27], which was also shown with MDCK-SIAT1 [21], MDCK-AX4 [22], and hCK [15] cells. A possible explanation for this discrepancy may lie in the glycoproteins on which *N*-glycans, the presumed glycan receptors for IAVs [40], are presented. It has been suggested previously that only specific sialylated glycoproteins can be used as a receptor for IAV [40, 42], such as the voltage-dependent Ca^2+^ channel Ca_v_1.2 [43], NKp44 [44, 45], NKp46 [45-47], epidermal growth factor (EGFR) [48], and nucleolin [49]. Although we demonstrated that glycans on B3GNT2/B4GALT1 knock-in cells contained a higher number of LacNAc repeating units, we have not determined on which glycoproteins the elongated glycans are present. Possibly, the glycans that are used as a receptor and are present on specific glycoproteins can be modified in their SIA linkage, as done in MDCK-SIAT1, MDCK-AX4, and hCK cells, but not in the number of LacNAc repeating units. This would explain why the infection efficiency could not be increased by the elongation of LacNAc repeating units on MDCK and hCK cells.

Nevertheless, not all recent H3N2 IAVs could be isolated efficiently in hCK cells [15], as also shown in Fig. 5D. Possibly, no viable virus particles were present in the patient samples from which we attempted to isolate virus. Alternatively, we may be overlooking an identified [43-49] or an unidentified glycoprotein that is not present (in high enough quantities) on hCK cells. Additionally, our *N*-glycan analysis did not allow for exact glycan structure determination, and therefore it is unclear how many consecutive LacNAc repeating units are present on the glycans. Furthermore, other types of glycans, such as phosphorylated [50] and sulfated glycans [2, 10, 51], possibly act as a receptor for IAV. Due to our sample preparation for the released glycan mass spectrometry analysis, we were unable to measure phosphorylated and sulfated glycans. To increase the isolation of recent H3N2 IAVs, it is of foremost importance to investigate the limiting factor in the infection efficiency of these viruses.

The overexpression of B3GNT2 and/or B4GALT1 is responsible for the elongation of glycans. Previously, overexpression of B4GALT1 was found to result in the elongation of glycans on CHO cells [31]. From the flow cytometry analysis, B4GALT1 appeared to be the limiting factor for the elongation of glycans in MDCK cells, while the elongation was limited by B3GNT2 in hCK cells. The glycan mass spectrometry results showed that the relative abundance of glycans with high numbers of LacNAc repeating units was only marginally increased by the overexpression of either or both B3GNT2 and B4GALT1, while the glycans on hCK cells were elongated by the overexpression of either B3GNT2 or B4GALT1. The glycans that are investigated in both methods are different since we look at all glycans (*N*-glycans, *O*-glycans, and glycolipids) in flow cytometry experiments and only *N*-glycans during glycan mass spectrometry. In both methods, the hCK-B3GNT2 were shown to have the highest relative abundance of elongated LacNAc repeating units. In hCK cells, sialyltransferase expression is severely modified by the overexpression of ST6GAL1 and the knockout of all ST3GAL enzymes. Both the heavily overexpressed ST6GAL1 and B3GNT2 in hCK cells use galactose as a substrate. The overexpression of B3GNT2 in hCK cells perhaps restores the balance between ST6GAL1 and B3GNT2, thereby allowing B3GNT2 to use the galactose as a substrate again for the elongation of glycans before sialylation takes place. On the other hand, in MDCK cells, the balance may be skewed even more by the overexpression of B3GNT2, leading to the low sialylation of glycans on these cells (Fig. S4B).

Our observations indicate that only few suitable glycan receptors are required for efficient infection. This is in line with our previous observations that a slight increase from 2.7% to 8.7% of sialylated glycans with at least three consecutive LacNAc repeating units on turkey erythrocytes allowed for the binding of contemporary H3N2 viruses [12]. Also in ferrets, an animal model that is often used to study human influenza viruses [38], the presence of glycans in the respiratory tract (lung, trachea, soft palate, nasal turbinate, nasal wash) was investigated. Elongated glycans were present solely as *N*-glycans, with a maximum of 9 LacNAc repeating units per glycan, but at most 0.17% of the detected glycans had at least three consecutive LacNAc repeating units terminating with SIA, which is required for H3N2 IAV binding [52]. Although the glycans in the human trachea have not been analyzed yet, data is available on other parts of the human airway system. Sensitive methods indicated the presence of extended *N*-glycans with up to 10 LacNAc repeating units in human lung tissue. However, at most 0.3% of the *N*-glycans were found to contain at least 3 consecutive LacNAc repeating units. The *N*-glycans in the bronchus and nasopharynx contained a lower number of LacNAc repeating units than in the lung [53]. Another study found *N*-glycans with up to 22 LacNAc repeating units in the lung. Even though the majority of the SIAs were found to be α2,6-linked instead of α2,3-linked, the α2,6-linked SIAs were mainly present on the shorter glycans [54], further supporting that only minor amounts of suitable glycan receptors are required for efficient infection.

## Material and Methods

### Cell culturing and preparation of cell lysates

Cells were cultured in DMEM (Gibco) with 10% FCS (S7524, Sigma) and 1% penicillin and streptomycin (Sigma). All hCK cells [15], knock-in and WT, were maintained with an additional 10 µg/ml blasticidin and 2 µg/ml puromycin in the medium. B3GNT2 and B4GALT1 knock-in cells were maintained in medium containing an additional 300 µg/ml Hygromycin B, a concentration that was determined to kill MDCK and hCK without Hygromycin B resistance genes. Detaching of the (knock-in) MDCK and hCK cells was always done using TrypLE Express Enzyme (12605010, Thermo Fisher Scientific).

Cell lysates were obtained after first washing cell monolayers once using D-PBS (D5837, Sigma). Cells were subsequently harvested after incubation at 37°C for 20 min with TrypLE Express Enzyme. The cell suspension was centrifuged for 5 min at 250 rcf. The cell pellets were lysed by the addition of RIPA lysis buffer (20-188, Merck Millipore) supplemented with protease inhibitor (A32965, Thermo Fisher Scientific), which was vortexed for 20 sec. The suspension was incubated on ice for 30 min, after which it was centrifuged at 16500 rcf in a fixed-angle centrifuge at 4°C, after which the supernatant was used as cell lysate.

### Cloning of lentiviral transfer plasmids

Plasmid pCF525-EF1a-Hygro-P2A-mCherry-lenti [55] was a gift from Jennifer Doudna (Addgene plasmid # 115796) and was used as the backbone for the transfer plasmid. Three transfer plasmids were constructed (pCF-B3GNT2, pCF-B4GALT1, and pCF-B3GNT2-B4GALT1). The region between the P2A and WPRE was removed and replaced by either the *B3GNT2* or *B4GALT1*. When the genes of both glycosyltransferases were cloned into the plasmid they were connected with a T2A self-cleaving peptide. The *B3GNT2* and *B4GALT1* genes were always proceeded by the signal sequence of the human GalT, which we copied from the EGFP-GalT plasmid (gift from Jennifer Lippincott-Schwartz, Addgene plasmid # 11929) [56]. The T2A self-cleaving peptide was amplified from plasmid tetO.Sox9.Puro [57], which was a gift from Henrik Ahlenius (Addgene plasmid # 117269). The *B3GNT2* and *B4GALT1* genes were amplified from plasmids B3GNT2-pGEn2-DES and B4GALT1-pGEn2-DES, which are a gift from Kelly Moremen and are available via http://glycoenzymes.ccrc.uga.edu/. All segments were amplified with an overhang, using the primers indicated in Table 1. Assembly of the plasmids was performed using Gibson assembly, after which they were sequenced to ensure correct amplification and assembly.

**Table 1.**
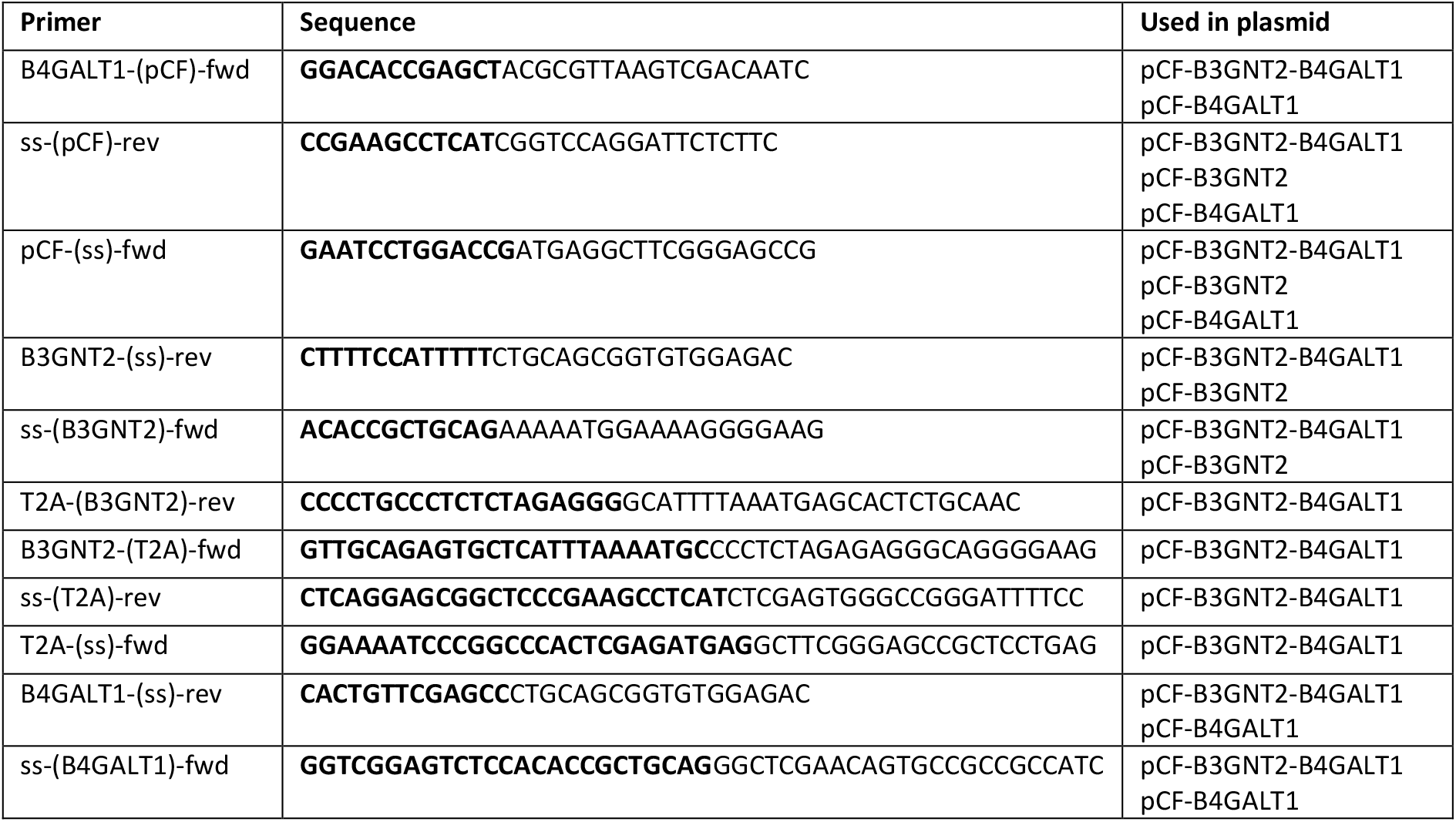
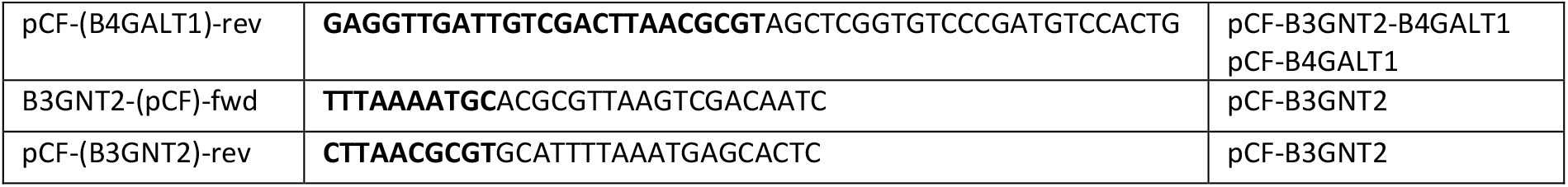
Primers used for the generation of the transfer plasmids. In brackets is indicated which segment is amplified, and the other part of the name indicates the overhang. The overhang is marked in bold in the sequence.

### Lentiviral integration of the *B3GNT2* and *B4GALT1* genes

Lentiviral particles were produced using HEK293T cells [58]. One of the transfer plasmids as described above, together with the packaging plasmids pMDLg/pRRE, pRSV-Rev, and pMD2.G were used, which were kind gifts from Didier Trono [59] (Addgene plasmids #12251, #12253, and #12259 respectively). The day before transduction, MDCK and hCK cells were seeded in a 6 wells plate at a density of 100.000 cells per well. Transduction with 0.5-3 µl of lentivirus was performed in presence of 8 µg/ml polybrene with 1 ml fresh medium per well. The medium was replaced with fresh medium containing 300 µg/ml Hygromycin B at 18 hours after transduction. Cells were grown until no Hygromycin B sensitive cells were remaining. Cells were always maintained in the presence of 300 µg/ml Hygromycin B.

### RT-qPCR analysis on *B3GNT2, B4GALT1*, and *ST6GAL1* genes

RNA extraction was performed using the GeneJET RNA purification kit (Thermo Fisher Scientific) according to the manufacturer’s protocol, after which the DNA was treated with DNAse I (#EN0251, Thermo Fisher Scientific). RT-qPCR was performed using the Luna universal one-step RT-qPCR kit (#E3005, New England Biolabs) according to the provided protocol, in which 10 ng of DNAse I-treated RNA was used. Primers (Table 2) for *B3GNT2, B4GALT1*, and *ST6GAL1* were designed to anneal both in the human and dog genome. Primers for GAPDH (household/reference gene) were designed using the dog genome. Experiments were performed in triplicate and Ct values of the RT-qPCR experiments on the glycosyltransferases were compared to the average Ct value of GAPDH of that specific cell line under the assumption that the amount of DNA doubles every cycle. The means and standard deviations of the amount of DNA relative to GAPDH were calculated.

**Table 2.**
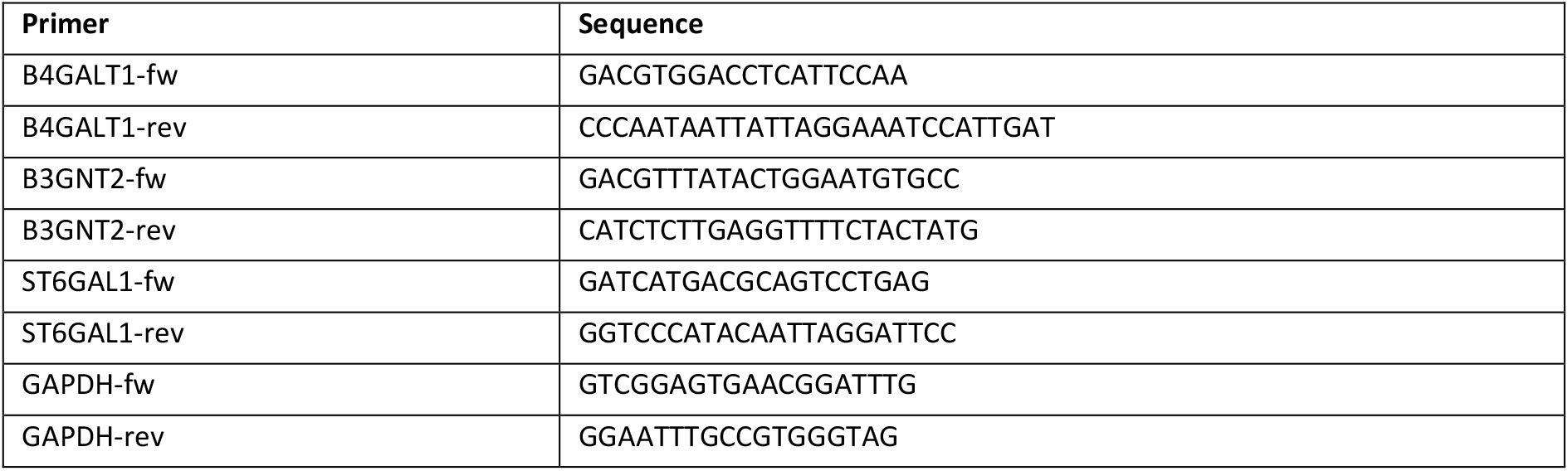
Primers used in the RT-qPCR experiments.

### Overexpression of B3GNT2, B4GALT1, and ST6GAL1 on the protein level

Cell lysates, obtained as described above, were further used for the label-free quantification of B3GNT2, B4GALT1, and ST6GAL1 proteins. From the cell lysates, 10 µg of protein was denatured, reduced, and alkylated by adding 100 µl of 150 mM Tris, 5mM TCEP (tris(2-carboxyethyl)phosphine), 30 mM chloroacetamide (CAA), 1% sodium deoxycholate (SDC) at pH 8.5. Next, 100 ng endoproteinase lysC and 100 ng trypsin were added and the samples were incubated overnight at 37°C. The samples were then acidified by adding formic acid (FA) to a concentration of 0.5% before solid-phase extraction (SPE) sample clean-up, causing the SDC to precipitate. SPE clean-up was performed on an Oasis HBL u-elution plate.

After the SPE clean-up, the samples were dried with a vacuum centrifuge. Subsequently, the sample was reconstituted in 2% FA before analysis on the Orbitrap Exploris mass spectrometer (Thermo Scientific) connected to a UHPLC 3000 system (Thermo). Approximately 200 ng of reconstituted peptides were trapped on a pre-column and then separated on a 50 cm x 75 μm Poroshell EC-C18 analytical column (2.7 μm) temperature controlled at 40°C. Solvent A consisted of 0.1% FA, solvent B of 0.1% FA in 80% acetonitrile, and different combinations of solvent A and B were used in the next steps. Trapping was performed for 2 min in 9% solvent B. Peptides were separated by a 65 min gradient of 9–44 % buffer B followed by 44–99% B in 3 min, and 99% B for 4 min. Mass spectrometry (MS) data were obtained in a data-dependent acquisition mode. The full scans were acquired in the m/z range of 350-1600 at the resolution of 60,000 (m/z 400) with AGC target 3E6. The most intense precursor ions were automatically selected for HCD fragmentation performed at normalized collision energy 28, after accumulation to the target value of 1E5. MS/MS acquisition was performed at a resolution of 15,000. Protein identification was done with Byonic (Protein Metrics). A search was performed against the dog proteome (UP000002254_9615) with the addition of the human B3GNT2, B4GALT1, and ST6GAL1 sequences. The search was performed with specific digestion C-terminal of R/K, allowing 3 missed cleavages, using precursor and fragment mass tolerances of 12 and 24 ppm, respectively. Carbamidomethylation of cysteine was set as a fixed modification and oxidation of the methionine or tryptophan as a variable modification. Peptides unique for the human B3GNT2, B4GALT1, and ST6GAL1 were manually selected and the MS1 peak areas were integrated with Skyline and normalized against the combined MS1 signals for identified peptides of tubulin beta (E2RFJ7). The peptide library for Skyline was built by repeating the search with a focused database containing only the human B3GNT2, B4GALT1, and ST6GAL1 and tubulin beta sequences. The mass spectrometry proteomic data have been deposited to the ProteomeXchange Consortium via the PRIDE [60] partner repository with the dataset identifier PXD037175.

### Expression and purification of trimeric HA for binding studies

Recombinant trimeric IAV hemagglutinin proteins (HA) were cloned into the pCD5 expression vector as described previously [61, 62], in frame with a GCN4 trimerization motif (KQIEDKIEEIESKQKKIENEIARIKK), a superfolder GFP [39] or mOrange2 [63] and the Twin-Strep-tag (WSHPQFEKGGGSGGGSWSHPQFEK); IBA, Germany). The open reading frames of the HAs of A/Vietnam/1203/2004 H5 (Addgene plasmid #182546, [35]), A/Puerto-Rico/8/1934 H1 [39], A/Hong-Kong/480/2014 H3 (3C.2a), A/Netherlands/109/2003 H3, A/Netherlands/761/2009 H3, A/Netherlands/354/2016 H3, A/Netherlands/00010/2019 H3 (3C.2a), A/Hong-Kong/1/1968 H3, A/Singapore/INFH-16-0019/2016 H3 (3C.2a), A/Kansas/14/2017 H3 (3C.3a), and the NTD of γCoV/AvCoV/guinea fowl/France/14032/2014 were synthesized and codon-optimized by GenScript. The trimeric HAs were expressed in HEK293S GnTI(-) cells with polyethyleneimine I (PEI) in a 1:8 ratio (µg DNA:µg PEI) for the HAs as previously described [61], while a 1:12 ratio was used for the NTD of the guinea fowl CoV. The transfection mix was replaced after 6 hours by 293 SFM II suspension medium (Invitrogen, 11686029), supplemented with sodium bicarbonate (3.7 g/L), Primatone RL-UF (3.0 g/L, Kerry, NY, USA), glucose (2.0 g/L), glutaMAX (1%, Gibco), valproic acid (0.4 g/L) and DMSO (1.5%). Culture supernatants were harvested 5 days post-transfection and purified with sepharose strep-tactin beads (IBA Life Sciences, Germany) according to the manufacturer’s instructions.

### Flow cytometry studies

Cells were harvested using TrypLE Express Enzyme as described above. After removal of the supernatant, cells were resuspended in PBS supplemented with 1% FCS (S7524, Sigma) and 2mM EDTA and kept at 4°C at any time. For experiments with α2-3,6,8,9 neuraminidase A (#P0722, New England Biolabs), neuraminidase (NA) was used 1:200 with 1,000,000 cells per ml in glycobuffer 1 (5 mM CaCl_2_, 50 mM sodium acetate, in MQ water, at pH 5.5) for 16 hours at 37°C on a shaking platform in the dark, before incubation with the lectin/HA mixes. In a round-bottom 96-wells plate (353910, Falcon), 150,000 cells were used. Per well, 100 µl of PBS supplemented with 1 µg of HA or biotinylated lectin (SNA (B1305), LEL (B1175), ECA (B1145), all from Vector Laboratories) was used, to achieve a final concentration of 10 µg/ml. Hemagglutinins were precomplexed (on ice, 20 min) with 1.3 µg monoclonal antibody detecting the Twin-Strep-tag and 0.325 µg goat anti-human Alexa Fluor 488 (A11013, Invitrogen). Biotinylated lectins were precomplexed (on ice, 20 min) with 0.2 µg streptavidin Alexa Fluor 488 (S32354, Invitrogen). For titration experiments, different amounts of HA, lectin, precomplexing antibodies, or streptavidin were used. Furthermore, eBioscience Fixable Viability Dye eFluor 780 (65-0865, Thermo Fisher Scientific) was diluted 1:2000 in the same mixture. Cells were incubated with the hemagglutinin/lectin mixed for 30 minutes at 4°C in the dark. Cells were washed once with PBS supplemented with 1% FCS and 2 mM EDTA, after which the cells were fixated with 100 µl of 1% paraformaldehyde in PBS for 10 minutes. Afterward, cells were washed once using PBS supplemented with 1% FCS and 2 mM EDTA, after which they were resuspended in 100 µl of the same buffer. Flow cytometry was performed using the BD FACSCanto II (BD Biosciences) using appropriate laser voltages. Data were analyzed using FlowLogic (Inivai Technologies) and gated as described in Fig. 2A to consecutively select cells, single cells, and cells that are not dead. Mean fluorescence values of triplicates were averaged and standard deviations were calculated. Curves for titration experiments were smoothed using the standard settings.

### Identification of *N*-glycans on cells by mass spectrometry

Cell lysates of WT and B3GNT2/B4GALT1 knock-in MDCK and hCK cells were obtained as described above. The total protein concentration in the cell lysates was determined using a BCA assay. The glycans in 400 µg of total protein were released by PNGaseF treatment. Proteins were first denatured in DTT/SDS (40 mM DTT, 0.5% v/v SDS) for 8 minutes at 95°C, after which they were cooled on ice. Subsequently, NP-40 (1% v/v) and glycobuffer G7 (50 mM sodium phosphate at pH 7.5) were added, together with 30 µg of PNGaseF. The samples were incubated in a shaking incubator overnight at 37°C. Samples were centrifuged (4700 rcf, 3 min) to remove potential precipitate, after which they were loaded on separate C18 SPE cartridges (Avantor™ 7020-02 BAKERBOND™ SPE Octadecyl), which were beforehand conditioned with 1 ml acetonitrile (MeCN) and 1 ml MQ water. The flow-through was collected and the remaining glycans were eluted from the C18 cartridges with 1 ml of 5% MeCN and 0.05% trifluoroacetic acid (TFA) in MQ water. The MeCN and TFA in both samples were evaporated under a stream of nitrogen gas. Flow through and elution fractions were diluted into 500 µl MQ water and combined, after which PGC SPE cartridges (Thermo Scientific™ HyperSep™ Hypercarb™ SPE cartridges) were used to further purify the samples. The PGC SPE cartridges were conditioned with 1 ml MeCN and 1 ml MQ water, after which the samples were loaded on the cartridges. The cartridges were washed with 1 ml 0.05% TFA in MQ water and 1 ml 5% MeCN with 0.05% TFA in MQ water. Samples were eluted with 50% MeCN and 0.1% TFA in MQ water and evaporated under a stream of nitrogen gas. The dried glycans were dissolved in 30 µl MQ water and 6 µl pure glacial acetic acid and labeled using 5 µl procainamide (105 mg/ml procainamide HCl in DMSO) and 5 µl 2-picoline borane (107 mg/ml 2-Methylpyridine borane complex in DMSO). The solution was vortexed thoroughly and incubated for 2 hours at 65°C, after which the samples were evaporated using the vacuum concentrator. The sample was dissolved in 300 µl MQ water and vortexed until the pellets were dissolved, after which 5 µl 25% (w/v) ammonia was added per sample to ensure a pH above 10. To remove the unused procainamide from the reaction mixture, liquid-liquid extraction with 500 µl dichloromethane was performed three times, with centrifuge steps of 4700 rcf for 3 min in between. The dichloromethane was removed and residual dichloromethane was evaporated under a stream of nitrogen gas. The samples were dissolved in a total of 1 ml MQ water after which they were loaded onto PGC SPE cartridges (conditioned with 2 ml MeCN and 2 ml MQ water). The cartridges were washed with 2 ml MQ water and the glycans were eluted using 50% MeCN with 0.1% TFA in MQ water, after which the MeCN and TFA were evaporated under a stream of nitrogen gas, followed by lyophilization.

Before HILIC-IMS-QTOF analysis, the lyophilized samples were reconstituted in 15 µl 70% MeCN in MQ water and centrifuged. The injected volume was 10 µl. The HILIC-IMS-QTOF system was an Agilent 1260 Infinity LC coupled to a 6560 IM-QTOF mass spectrometer (Agilent Technologies, Santa Clara, USA). For HILIC separation, a SeQuant ZIC-cHILIC column (3 µm, 100 Å; 150 × 2.1 mm) was used with a matching guard column (20 × 2.1 mm). The temperature of the column compartment was set at 40 °C. The mobile phase was composed of eluent A: 10 mM ammonium formate with 10 mM formic acid in MQ water, and eluent B: LC-MS grade MeCN. The initial eluent composition was 30% A at a flow rate of 0.2 ml/min, followed by a linear gradient to 50% A from 0 to 20 minutes. 50% A was held isocratically until 25 minutes. To re-establish initial conditions, the column was flushed with at least 10 column volumes of 30% A.

The IMS-QTOF was set to positive ion mode with a capillary voltage of 3500 V, nozzle voltage of 2000 V, and a fragmentor voltage of 360 V. The drying gas temperature was 300 °C with a flow rate of 8 l/min and the sheath gas temperature was 300 °C at 11 l/min. The nebulizer pressure was set at 40 psi. The ion mobility settings were set as follows: 18 IM transients per frame, an IM trap fill time of 3900 µs and a release time of 250 µs, the drift tube voltage was 1400 V, and the multiplexing pulsing sequence length was 4 bits.

IM-MS data was calibrated to reference signals of m/z 121.050873 and 922.009798 using the IM-MS reprocessor utility of the Agilent Masshunter software. The mass-calibrated data was then demultiplexed using the PNNL preprocessor software using a 5-point moving average smoothing and interpolation of 3 drift bins. To find potential glycan hits in the processed data, the ‘find features’ (IMFE) option of the Agilent IM-MS browser was used with the following criteria: ‘Glycans’ isotope model, limited charge state to 5 and an ion intensity above 500. The found features were filtered by m/z range of 300 – 3200 and an abundance of over 500 (a.u.) where abundance for a feature was defined as ‘max ion volume’ (the peak area of the most abundant ion for that feature).

After exporting the list of filtered features, glycans with a mass below 1129 Da (the mass of an *N*-glycan core) were removed. The ExPASy GlycoMod tool [64] was used to search for glycan structures (monoisotopic mass values, 5 ppm mass tolerance, neutral, derivatized N-linked oligosaccharides, procainamide (mass 235.168462302) as reducing terminal derivative, looking for underivatized monosaccharide residues (Hexose, HexNAc, Deoxyhexose, and NeuAc)). For features with multiple potential monosaccharide combinations, the most realistic glycan in the biological context was chosen. The abundance of glycan features with the same mass, composition, and a maximum difference of 0.1 min in the retention time were combined as one isomer. Full glycan composition feature lists for the different cell lines are presented in Table S1-8.

Analysis of the number of LacNAc repeating units was performed on the complex and hybrid *N*-glycans with at least one LacNAc repeating unit. A glycan with one LacNAc repeating unit was defined as a glycan with 4 hexoses and a minimum of 3 HexNAcs or 3 HexNAcs and at least 4 hexoses. A glycan with two LacNAc repeating units was defined as a glycan with 5 hexoses and a minimum of 4 HexNAcs or 4 HexNAcs and at least 5 hexoses. This pattern was continued for the higher numbers of LacNAc repeating units. The total absolute abundance of all selected glycans was added up, after which the relative abundance of a given number of LacNAc repeating units was calculated from this total. Additionally, the percentage of these glycans with at least one SIA was calculated.

Chromatograms of the *N*-glycans with two to seven LacNAc repeating units, calculated as described above, from hCK WT and hCK-B3GNT2 cells were constructed using Agilent’s Masshunter Qualtitative Analysis 10.0 software (Fig. 3A). The shown chromatograms are the summed extracted-ion-count (EIC) for the ten most abundant glycan features per LacNAc repeating unit group. The EIC for a glycan was set as the observed m/z value with a symmetrical 10 ppm expansion. Different ionization states of the same glycan that were found as a separate feature by the feature-finding software were also included in the summed EIC chromatogram.

### Sugar nucleotide analysis

Cells were grown to 60-70% confluency in a 6-wells plate, after which the medium was removed and the cells were washed twice with wash buffer (75 mM ammonium carbonate in MQ water, pH 7.4 (corrected with glacial acetic acid), at 4°C). The cells were then treated with 700 µl of extraction buffer (40% acetonitrile, 40% methanol, 20% MQ water, at 4°C) per well for 2 minutes, after which the supernatant was transferred to a vial. This extraction step is repeated for 3 minutes, after which the two extracts were pooled and centrifuged at 18000 rcf for 3 min. The supernatant was taken and dried in the vacuum concentrator. Samples were frozen at −80°C until analysis using an ion-pair UHPLC-QqQ 1290-6490 Agilent mass spectrometer by Glycomscan BV (Oss, the Netherlands) [41].

### Virus titration on B3GNT2/B4GALT1 knock-in MDCK and hCK cells

Virus titers in the virus stocks in Table 3 were determined using end-point titration in MDCK cells and inoculated cell cultures were tested for agglutination activity using turkey red blood cells as an indicator of virus replication in the cells. For recent (2017-2019) H3N2 viruses, no binding to erythrocytes was observed and therefore virus titers were determined using a nucleoprotein (NP) staining. The NP staining was performed on the inoculated cells that were fixed with acetone for at least 20 minutes at −20°C. Primary mouse anti-NP antibody (HB65, 2 mg/ml) was diluted 1:3000 and the secondary goat anti-mouse IgG HRP antibody (A16702, 1 mg/ml, Thermo Fisher Scientific) was used at a dilution of 1:30000, after which 50 µl per well was used for both solutions. True Blue substrate (KPL) was then added to visualize positive wells using an ImmunoSpot Analyzer (CTL Europe, Bonn, Germany). Based on the negative control values and the highest positive values per plate, the cut-off for positivity was determined. Infectious titers were calculated from five replicates using the Spearman-Kärber method [65].

**Table 3.**
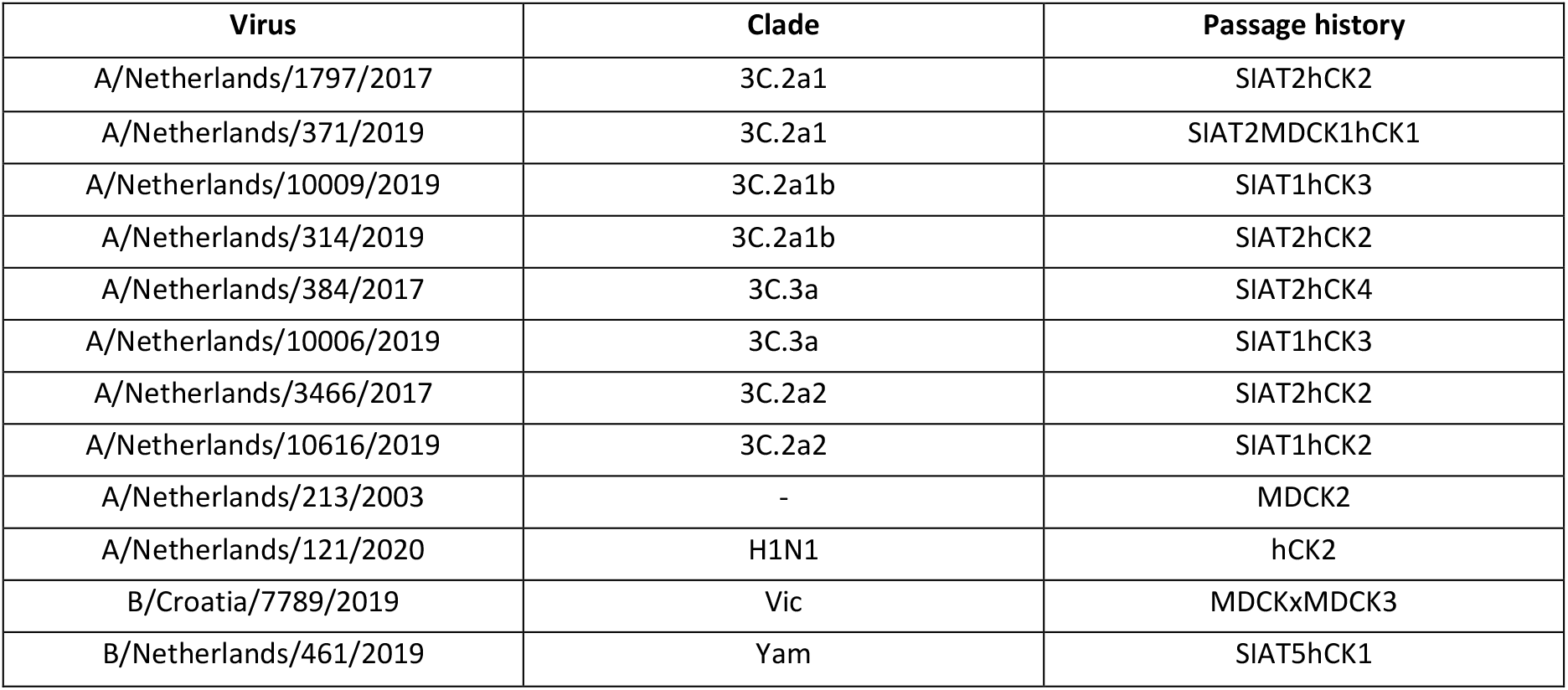
Details of IAVs used in the experiment shown in Fig. 5A-C. These viruses include one older H3N2 virus from 2003, one H1N1 virus, and two influenza B viruses. The viruses were passaged in MDCK, MDCK-SIAT1 [21], and/or hCK cells.

### Inoculation of hCK and hCK-B3GNT2 cells with influenza viruses

To evaluate whether IAVs that could not be isolated previously in hCK cells would replicate in hCK-B3GNT2 cells, hCK and hCK-B3GNT2 cells were seeded at a density of 20.000 cells per well in 96 wells plates at 24 hours before inoculation. The original patient material (100 µl) containing influenza virus (details of viruses in Table 4) was diluted in 700 µl infection medium (EMEM (Cambrex, Heerhugowaard, The Netherlands) supplemented with 100 U/ml penicillin, 100 μg/ml streptomycin, 2mM glutamine, 1.5mg/ml sodium bicarbonate (Cambrex), 10mM Hepes (Cambrex), nonessential amino acids (MP Biomedicals) and 20 μg/ml trypsin (Cambrex)), after which a two-fold dilution series was made. After three days, the cytopathic effects in the cells were evaluated and the mean (n=3) of the number of infected wells was calculated.

**Table 4.** Details of IAVs used in the experiment shown in Fig. 5D. The exact virus is indicated as well as the clade and HA mutations if applicable. Previous attempts of isolating these viruses in hCK cells were either unsuccessful (-) or successful (+).

| Number | Virus | Clade (+ HA mutations) | Cultured previously |
| --- | --- | --- | --- |
| 1 | A/Netherlands/3425/2017 | 3C.2a | - |
| 2 | A/Netherlands/2362/2018 | 3C.2a1b | - |
| 3 | A/Netherlands/2380/2018 | 3C.2a1b | - |
| 4 | A/Netherlands/010/2019 | 3C.2a1b | - |
| 5 | A/Netherlands/173/2019 | 3C.2a1b | - |
| 6 | A/Netherlands/1268/2019 | 3C.2a4 | - |
| 7 | A/Netherlands/1735/2019 | 3C.2a1b + T135K | - |
| 8 | A/Netherlands/1747/2019 | 3C.2a1b + T135K | - |
| 9 | A/Netherlands/1439/2019 | 3C.2a1b | + |
| 10 | A/Netherlands/1734/2019 | 3C.2a1b + T135K | + |
| 11 | A/Netherlands/054/2020 | 3C.2a1b + T131K | + |
| 12 | A/Netherlands/008/2021 | 3C.2a1b.2a2 | + |

## Acknowledgments

We thank professor Yoshihiro Kawaoka for providing the hCK cells. We would like to thank Frederik Broszeit for producing the glycans that are used in the glycan microarray in Fig. S1. Balthasar Heesters is thanked for his advice on the flow cytometry experiments and analysis. Monique van Scherpenzeel is thanked for the sugar nucleotide analysis. We also thank Roosmarijn van der Woude for her technical assistance.

## Supplementary figures

**Fig S1.**
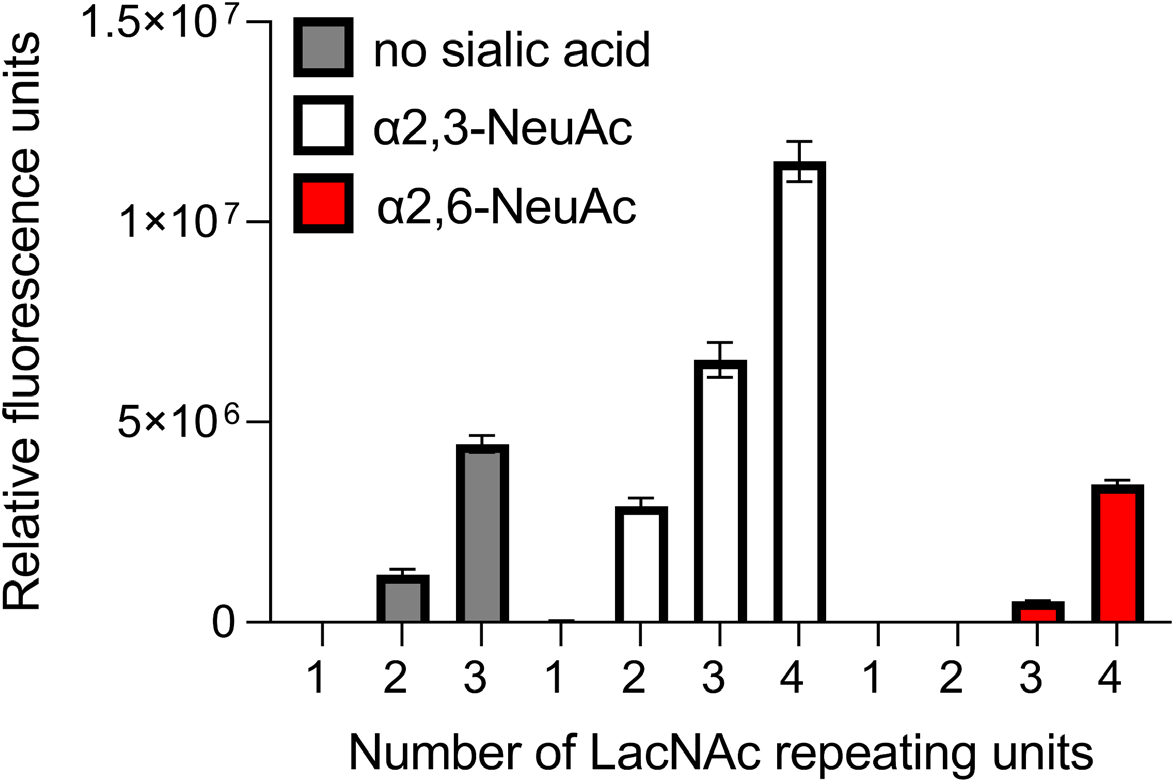
Binding specificity of *Lycopersicon esculentum* lectin on the glycan microarray. The binding of the *Lycopersicon esculentum* lectin (LEL) to symmetric bi-antennary *N*-glycans with 1, 2, 3, or 4 consecutive LacNAc repeating units terminating in no sialic acid, α2,3-linked NeuAc, or α2,6-linked NeuAc was investigated. Six replicates were performed simultaneously, after which the highest and lowest replicates were removed, and the mean and standard deviation were calculated over the four remaining replicates. The glycan array was performed as described earlier [35].

**Fig S2.**
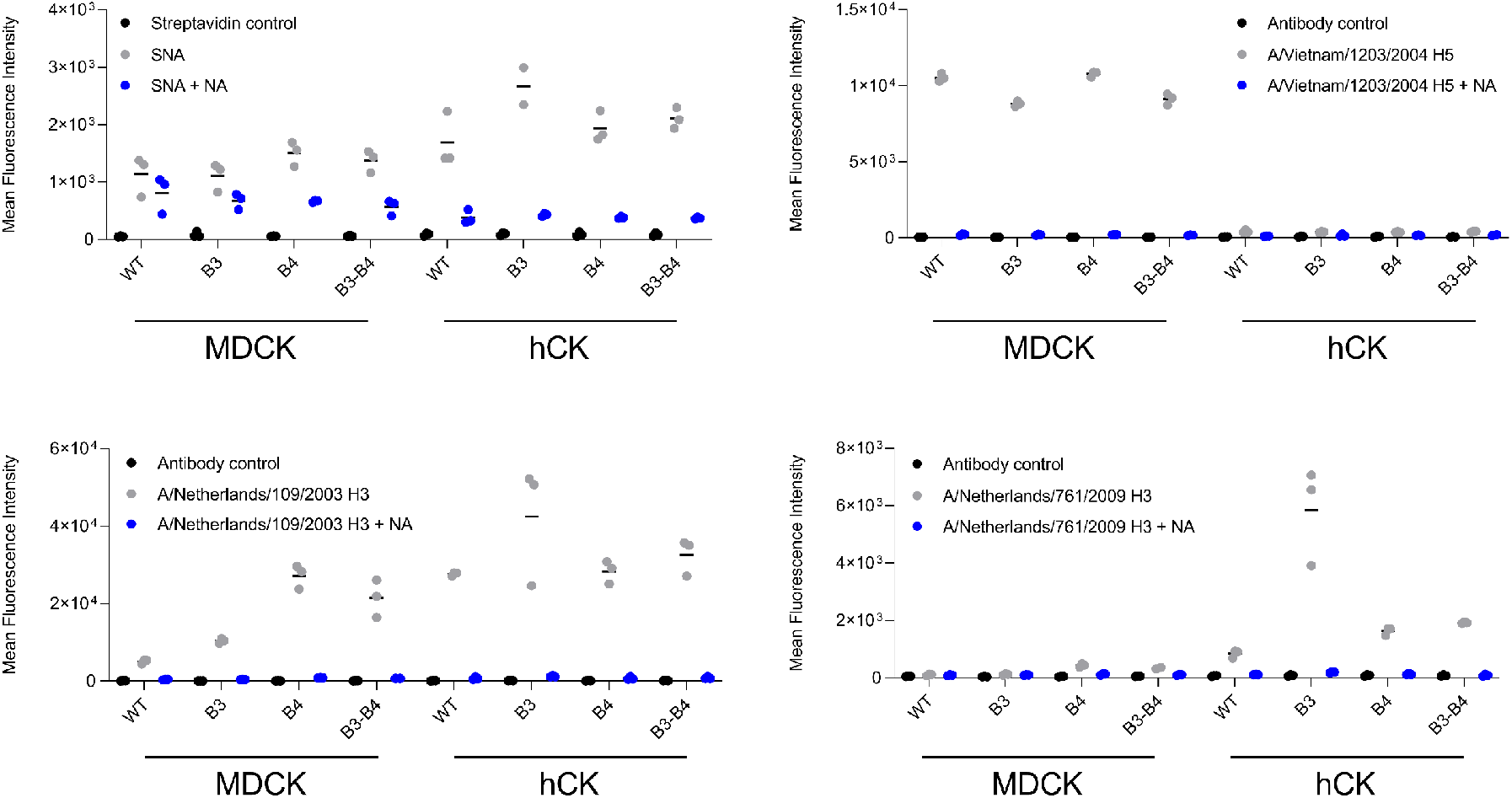
Flow cytometry with neuraminidase-treated B3GNT2/B4GALT1 knock-in MDCK and hCK cells. Binding of the lectin SNA and HAs A/Vietnam/1203/2004 H5, A/Netherlands/109/2003 H3, and A/Netherlands/761/2009 H3 with and without neuraminidase (NA) were measured using flow cytometry. The gating strategy as indicated in Fig. 2A was used. Triplicate measurements were performed and the mean and all individual measurements are shown.

**Fig S3.**
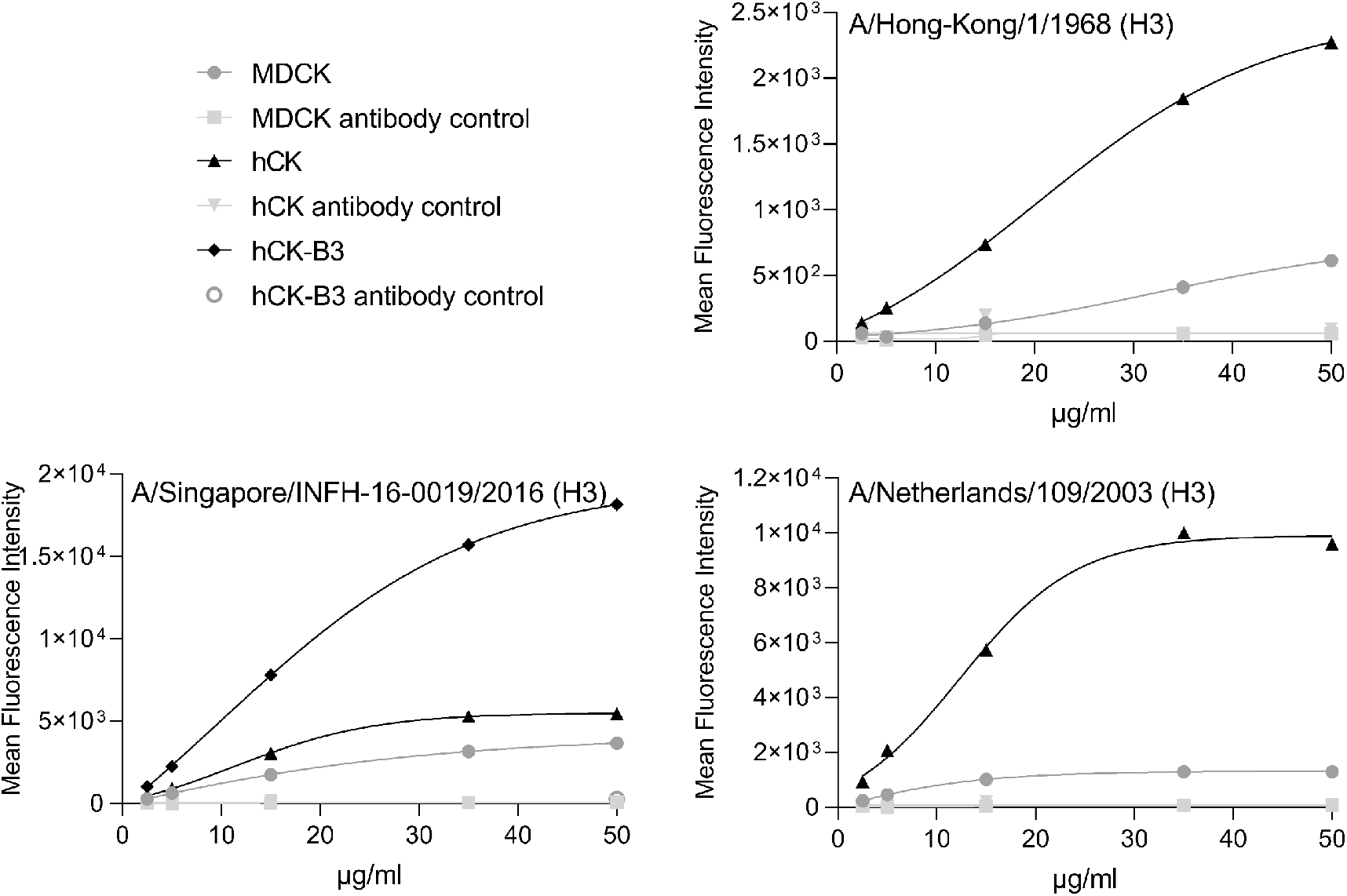
Titration of influenza hemagglutinins in flow cytometry. Flow cytometric titrations of the H3 HAs of A/Hong-Kong/1/1968, A/Singapore/INFH-16-0019/2016, and A/Netherlands/109/2003 in MDCK and hCK cells, including controls in the presence of just precomplexing controls were performed. For A/Singapore/INFH-16-0019/2016, also hCK-B3GNT2 cells were used. The gating strategy as described in Fig. 2A was used.

**Fig S4.**
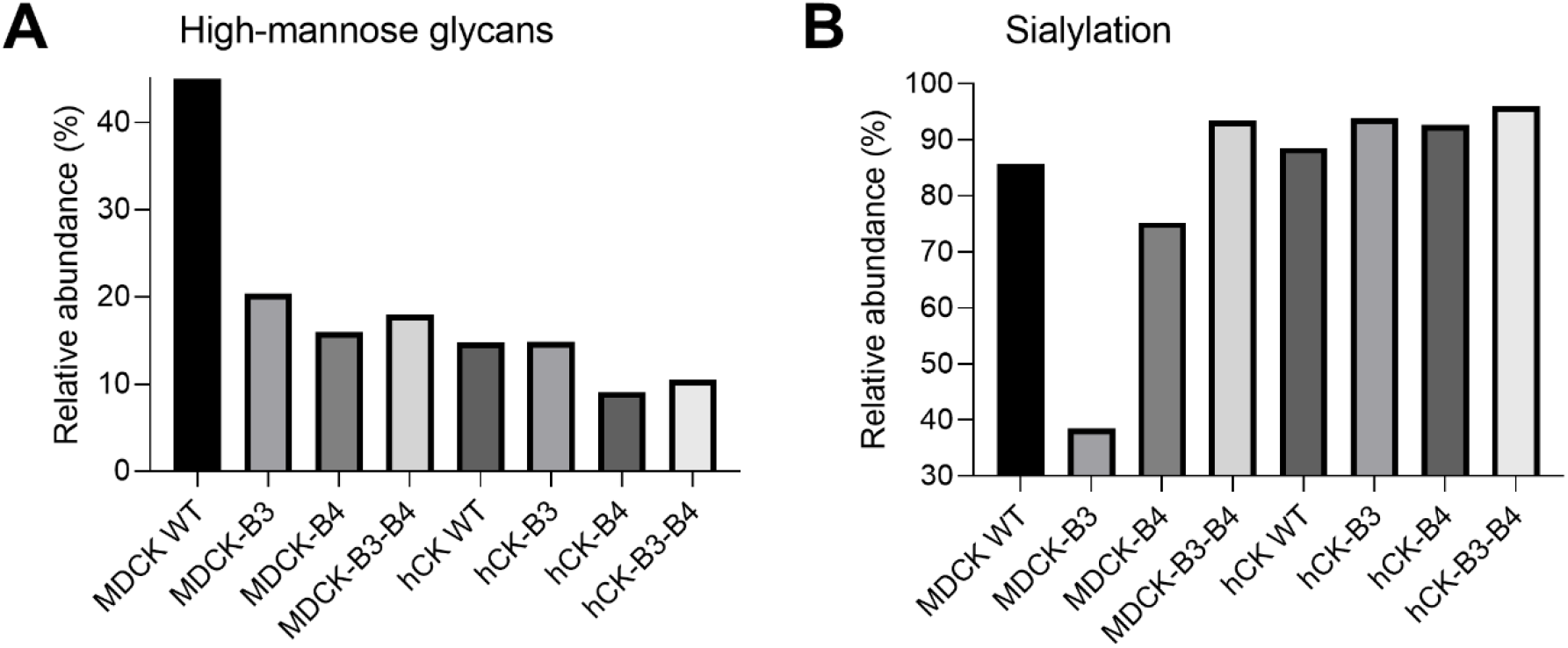
Relative abundance of high-mannose glycans and sialylation on WT and B3GNT2/B4GALT1 knock-in MDCK and hCK cells. The *N*-glycans from WT and B3GNT2/B4GALT1 knock-in MDCK and hCK cells were measured using mass spectrometry. (**A**) The relative abundance of high-mannose glycans was calculated as a percentage of all detected *N*-glycans (see Table S1-8). (**B**) The relative abundance of glycans (30-100%) with at least one SIA was calculated as a percentage of the total abundance of glycans with at least one LacNAc repeating unit (the glycans shown in Fig. 3B).

**Fig S5.**
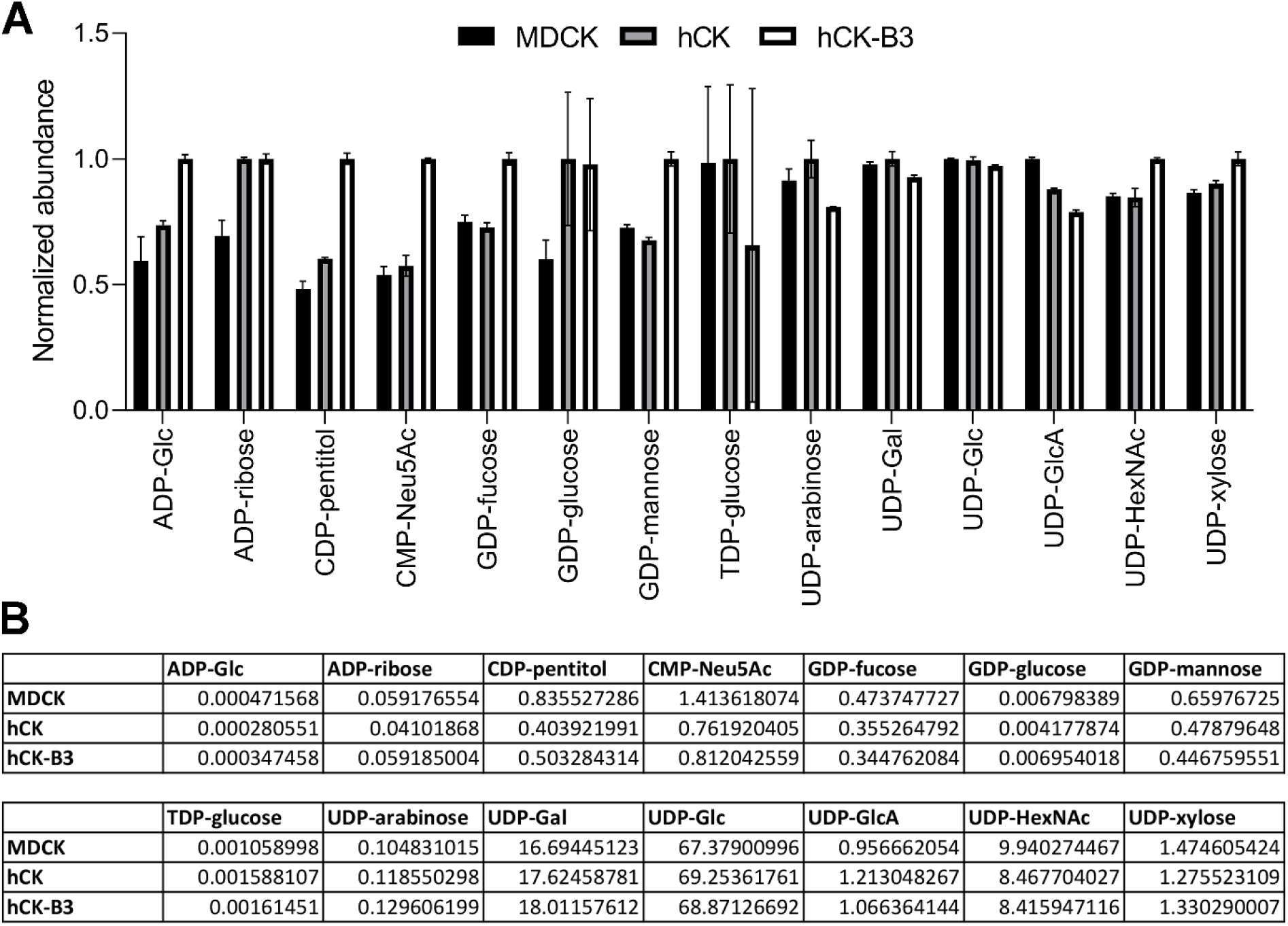
Sugar nucleotide analysis of MDCK, hCK, and hCK-B3GNT2 cells. The sugar nucleotides in the lysate of MDCK, hCK, and hCK-B3GNT2 cells were analyzed by mass spectrometry (n=2). (**A**) The normalized abundance of all measured sugar nucleotides is shown. Normalization was performed on the cell line with the highest amount of each sugar nucleotide. (**B**) Details of all analyzed sugar nucleotides, normalized over the sum of all measured nucleotide sugars.

